# Profiling of the reprogramome and quantification of fibroblast reprogramming to pluripotency

**DOI:** 10.1101/2020.02.25.964296

**Authors:** Kejin Hu, Lara Ianov, David Crossman

## Abstract

Pluripotent state can be established via reprogramming of somatic nuclei by factors within an oocyte or by ectopic expression of a few transgenes. Considered as being extensive and intensive, the full complement of genes to be reprogrammed, however, has never been defined, nor has the degree of reprogramming been determined quantitatively. Here, we propose a new concept of reprogramome, which is defined as the full complement of genes that need to be reprogrammed to the expression levels found in pluripotent stem cells (PSCs). This concept in combination with RNA-seq enables us to precisely profile reprogramome and sub-reprogramomes, and study the reprogramming process with the help of other available tools such as GO analyses. With reprogramming of human fibroblasts into PSCs as an example, we have defined the full complement of the human fibroblast-to-PSC reprogramome. Furthermore, our analyses of the reprogramome revealed that WNT pathways and genes with roles in cellular morphogenesis have to be extensively and intensely reprogrammed for the establishment of pluripotency. We further developed the first mathematical model to quantitate the overall reprogramming, as well as reprogramming in a specific cellular feature such as WNT signaling pathways and genes regulating cellular morphogenesis. We anticipate that our concept and mathematical model may be applied to study and quantitate other reprogramming (pluripotency reprogramming from other somatic cells, and lineage reprogramming), as well as transcriptional and epigenetic differences between any two types of cells including cancer cells and their normal counterparts.

## Introduction

An enucleated oocyte can reprogram an implanted somatic cell nucleus to pluripotent stem cells (PSCs) [1, 2]. Ectopic expression of a few transgenes can also induce pluripotent stem cells (iPSCs) from somatic cells, most commonly fibroblasts [3, 4]. iPSC reprogramming from human fibroblasts is a prolonged and stochastic process with a very low efficiency [4–6]. One reason for this inefficient conversion of cell fates is probably the great expanse of reprogramming required [7, 8]. Although it is considered to be extensive as well as intensive, the degree of iPSC reprogramming has not been determined quantitatively. A method for the measurement of reprogramming expanse is yet to be developed. Here, we report a new concept, reprogramome, which provides a basis for measurement of reprogramming. Subsequently, we developed the related concepts of downreprogramome, upreprogramome, erasome, and activatome. Using these concepts, we have precisely defined the breadth of reprogramming required for the establishment of human pluripotency. We have additionally developed mathematical models for quantification of reprogramming intensity of each gene in reprogramming and the total expanse of reprogramming using a new reprogramming unit, log2-transformed fold changes (LFC).

## Results

### Definition of reprogramome

We define reprogramome as the subset of genes that have to be reprogrammed so that one cell type can be converted into another one. A reprogramome generally includes two subgroups, downreprogramome and upreprogramome. Downreprogramome refers to the group of genes whose expression levels have to be downregulated while upreprogramome include the group of genes whose expression levels need to be upregulated for a complete conversion of cell fates. A downreprogramome may include a subset of genes whose expression has to be shut off completely, i.e., erasome. On the other hand, an upreprogramome may contain a subset of genes whose expression has to be activated *de novo*, with a term of activatome in its own right. Reprogramome may be a concept of transcription or epigenetics, and therefore there are transcriptional reprogramome and epireprogramome, respectively. Transcriptional reprogramome is a special sub-transcriptome, while epireprogramome is a defined sub-epigenome. Reprogramome may characterize any conversion of cell fates including pluripotency and lineage reprogramming. As a proof of principle, below we will summarize our profiling of the human transcriptional reprogramome for fibroblast conversion to iPSCs.

In the case of human fibroblast reprograming to iPSCs, the transcriptional downreprogramome should be the group of genes that have higher expression levels in fibroblasts than in PSCs. This group of genes has to be downregulated to the expression levels found in PSCs. On the other hand, the transcriptional upreprogramome is the group of genes that have higher expression levels in PSCs than in fibroblasts. These genes have to be upregulated to the levels found in PSCs. Therefore, in order to define the downreprogramome and upreprogramome, we just need to define the group of genes with higher expression in fibroblasts (fibroblast-specific genes, or simply fibroblast genes hereafter, or downregulatome) and the other group of genes with higher expression in PSCs (PSC genes hereafter, or upregulatome). The sum of the fibroblast-specific and PSC-specific genes constitutes the entire reprogramome of fibroblast-to-iPSC reprogramming.

### Extensive reprogramming revealed by reprogramome profiling

To this end, we sequenced RNA on human fibroblasts and PSCs. We used the NIH-registered human embryonic stem cell lines (ESCs), H1 and H9 because these are the widely used reference cell lines for PSCs [9]. Our RNA-seq is of high quality based on the quality control analyses and read counts for the signature genes of both cell types (see STAR Methods, and table S1). Using a set of strict criteria for selection (see STAR Methods), we showed that the downregulatome contains 3,617 genes/transcripts (Fig. 1A, Table S2), representing 26.4% of fibroblast transcriptome (Fig. 1G, Table S3). The upregulatome includes 4,190 genes/transcripts (Fig. 1B), equivalent to 30.6% of fibroblast transcriptome (Fig. 1G, and Table S3) and representing 28.8% of the ESC transcriptome (Table S3). Combining downregulatome and upregulatome, the reprogramome contains 7,807 genes/transcripts. This size of reprogramome is surprisingly large and is equivalent to 57% of the fibroblast transcriptome, and 53.6% of the ESC transcriptome. The actual reprogramome may be greater because our selection criteria may have excluded a subset of genes with very low expression levels, as well as the subset of genes whose differences in expression levels between the two types of cells are lower than 2 fold, the threshold we used.

**Fig. 1.**
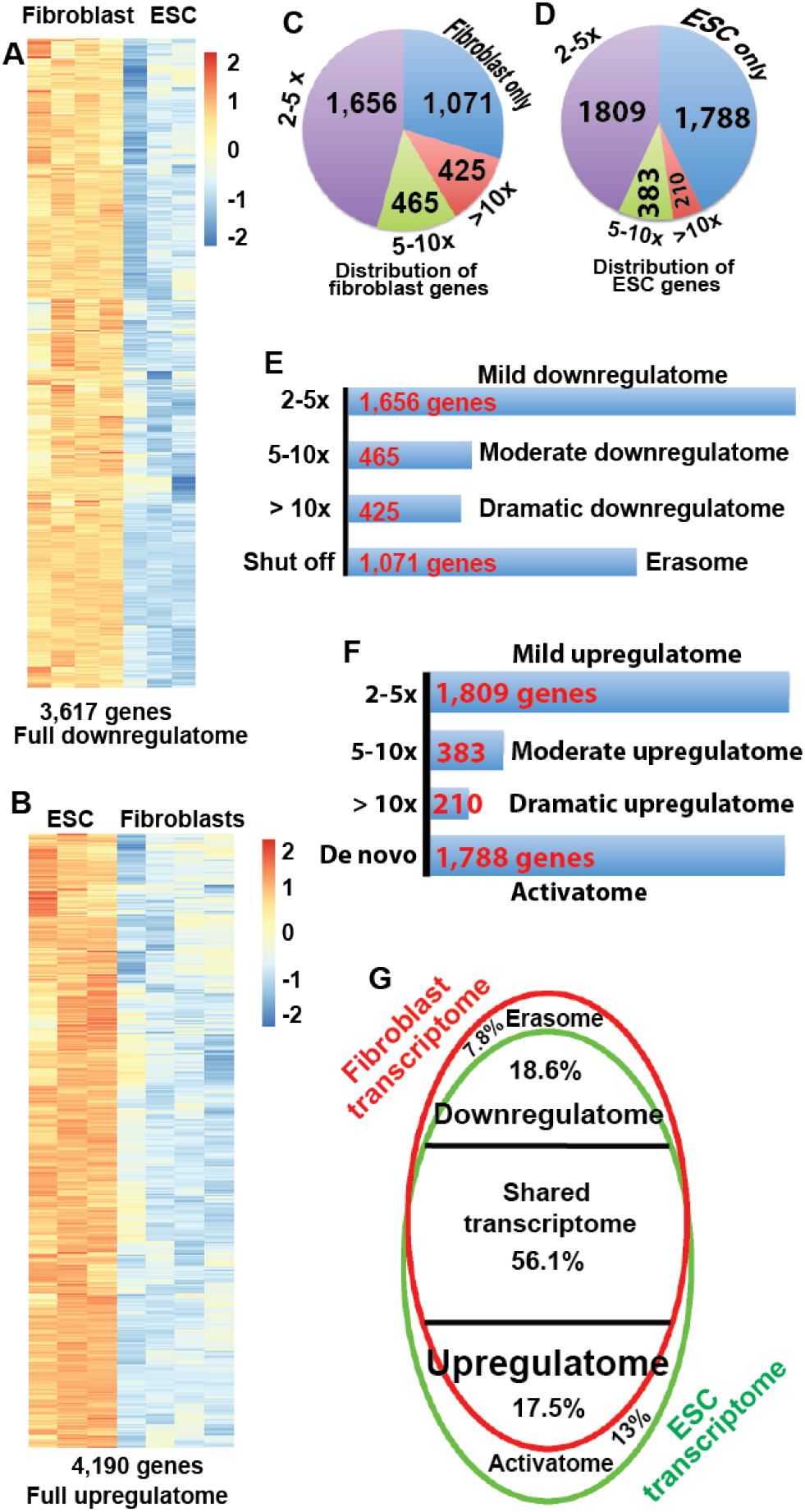
Profiling human fibroblast-to-iPSC reprogramome by RNA-seq. (A) Heat maps showing differences of gene expression levels with 2 fold or higher expression in human fibroblasts. Log2 scale. Fibroblast, n=4; ESC, n=3. q<0.01. (B) Heat maps showing differences of gene expression levels with 2 fold or higher expression in human ESCs. Log2 scale. Fibroblast, n=4; ESC, n=3. q<0.01. (C) Distributions of differentially expressed genes into different levels of fold changes for fibroblast. (D) Distributions of differentially expressed genes into different levels of fold changes for ESC genes. (E) Numbers of genes in different groups of reprogramming levels for dowregulatome. (F) Numbers of genes in different groups of reprogramming levels for upregulatome. (G) Relative size to fibroblast transcriptome for the different sub-reprogramomes.

### iPSC reprogramming is intensive

Next, we investigated the intensity of reprogramming. To this end, we broke the differences in gene expression levels between fibroblasts and ESCs into four tiers: 1) expressed in one cell type only but not in the other one; and differences in gene expression levels between the two cell types is 2) greater than 10 fold changes (FC), 3) 5 to 10 FC, and 4) 2 to 5 FC (Figs. 1C-F). The first tier includes activatome and erasome as defined above, representing the most radical reprogramming. The remaining three tiers are designated as dramatic, moderate, and mild reprogramming. Our data show that the activatome includes 1,788 genes/transcripts equivalent to 13.1% of fibroblast transcriptome (Figs. 1D, 1F, 1G, and Tables S2 and S3), while the erasome contains 1,071 genes/transcripts, representing 7.8% of fibroblast transcriptome (Figs. 1C, 1E, 1G, and Table S2 and S3). Combining these two groups, the radical reprogramming tier includes 2,859 genes/transcripts, equivalent to 20.9% of the fibroblast transcriptome. There are 425 genes/transcripts in the category of dramatic downregulatome, and 210 genes/transcripts in dramatic upregulatome. The dramatic tier therefore includes 635 genes/transcripts, representing 4.6% of the fibroblast transcriptome. Thus, 3,494 genes/transcripts have to be reprogrammed dramatically (10 fold above) or radically, equivalent to 24% of the ESC transcriptome and 25.5% of the fibroblast transcriptome. From these data, it is evident that pluripotency reprogramming is both extensive and intensive involving 57% the size of fibroblast transcriptome and a large activatome and erasome.

As expected, some well-known pluripotent genes [10] are among the activatome, for examples, *CLDN6, DPPA4, GDF3, L1TD1, LEFTY1, LEFTY2, LIN28A, LRRN1, NANOG, NODAL, PRDM14, SALL3, SOX2, TDGF1, TERT, ZFP42, ZIC5,* and *ZSCAN10* (Table S2). Unexpectedly, the master pluripotent gene, OCT4A (POU5F1), is not in the list of activatome, but is in the dramatic upreprogramome. This is because we used FGF2 in our culture of fibroblasts and FGF2 has been reported to stimulate *OCT4* expression [11]. *OCT4* has a mean read counts of 124 in our fibroblasts, which is above the cutoff of 50. Some other established pluripotency genes are among the dramatic upregulatome, for examples, *DNMT3B, SALL2*, and *SALL4* (Table S2). *PODXL,* the gene encoding a carrier protein for the two widely used pluripotency surface markers TRA-1-60 and TRA-181 [12], is within the dramatic reprogramome. MYC, one of the original reprogramming factors [3, 5, 13], is a member of the moderate upregulatome. Interestingly, *KLF2, KLF4* and *KLF5,* the well-known pluripotency genes in mouse [14], are all among the human downregulatome rather than upregulatome, with *KLF2* unexpectedly in the erasome (Table S2). Although KLF4 is one of the four canonical reprogramming factors [3, 5, 13] and plays a role in human pluripotency [15], it is widely expressed in various types of cells and tissues [16]. Of note, *KLF4* was first cloned from fibroblasts [17].

### Extensive reprogramming in WNT pathway

To understand the unique features that have to be established during reprogramming, we further conducted gene ontology (GO) analyses with the activatome using the PANTHER platform [18]. Out of the 1,549 uniquely mapped genes, 1,405 genes fall into the unclassified group. Nevertheless, 275 genes in the activatome can be assigned to at least one PANTHER pathway. Out of the 163 pathways available in PANTHER databases, 100 are represented in the activatome, and 76 pathways are over-represented (Table S4). Among them, 29 pathways have a p value less than 0.05 and 19 pathways have FDR less than 0.05 (Table S4). Fig. S1A shows the top 20 pathways over-represented by activatome in terms of p values. Of note, 47 genes in WNT signaling pathway are in the group of activatome (p = 2.58 × 10^−5^) (Fig. 2A, Tables S4 and S5). Other interesting pathways include cadherin signaling (33 genes, p = 1.3 × 10^−6^), FGF signaling (18 genes, p = 0.01), and VEGF signaling (12 genes, p = 0.01). Considering that a large number of genes in WNT pathway are represented by the activatome, we further analyzed the WNT-pathway genes in the upreprogramome. Surprisingly, the upreprogramome includes 87 WNT-pathway genes (p = 5.62 × 10^−4^) (Fig. 2C, and Table S5). This prompted our further pathway analyses with erasome and found that it contains 19 WNT-pathway genes (Fig. 2B, and Table S5). We then analyzed the full downreprogramome and found that it contains 56 WNT-pathway genes (Fig. 2C, and Table S5). Therefore, genes in WNT pathways should be intensely and extensively reprogrammed (also see quantification below).

**Fig. 2.**
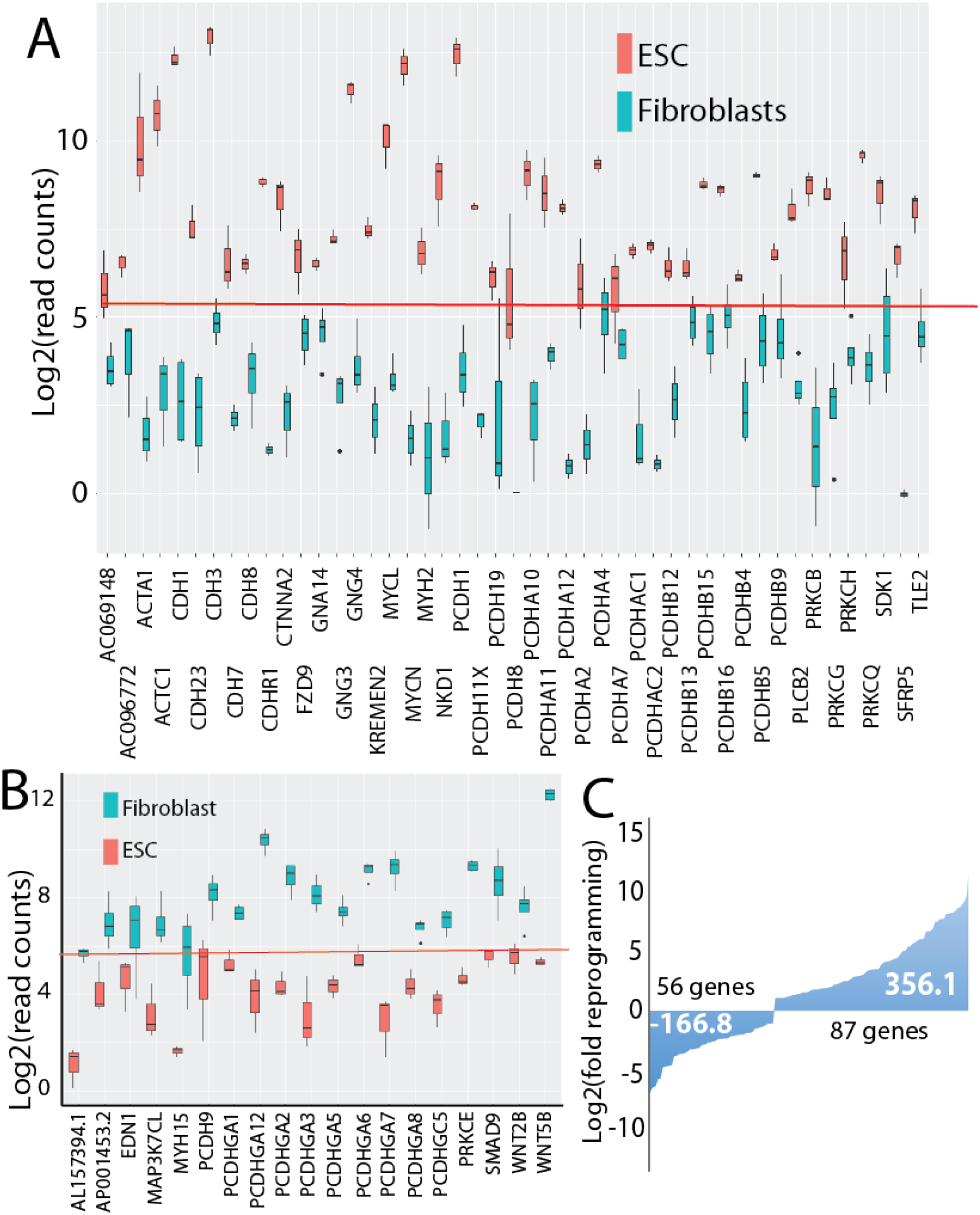
Profiling of WNT sub-reprogramome, and quantification of reprogramming in WNT pathways. (A) Box plot showing WNT genes that have to be activated de novo for iPSC generation. Red lines in A and B mark the threshold for expressed genes. (B) Box plot showing WNT genes that have to be shut off for iPSC generation. (C) Total reprogramming for genes in WNT pathways. Unit for y-axis is LFC. The number within the waterfall plot is the reprogramming values in LFC calculated using our mathematical model. Numbers of WNT genes to be down- and up-reprogrammed are indicated along the waterfall plots.

### A mathematic model for quantification of reprogramming

Pluripotency reprogramming is intensive and extensive, but there is no available method to quantitate reprogramming. After establishment of the reprogramome concept that allows for profiling of reprogramming genes, we further reasoned that the total expanse of pluripotency reprogramming could be measured by total numbers of genes to be reprogrammed along with the degree of reprogramming for each gene. The degree of reprogramming for each gene is reflected by its fold change (FC) in transcription. To distinguish down-from up-reprogramming, we propose to use the log2-transformed fold change (LFC). For gene *i*, the log2-transformed fold change to achieve for complete reprogramming is G*i* = log2(FC*i*) (Fig. 3A). We assume that under an ideal condition (for example, reprogramming that happens in a fertilized egg, or reprogramming in a reconstructed egg with a transferred somatic nucleus into a mature oocyte [19]), every gene has the same reprogramming constant. Given a reprogramming constant of α, the amount (or intensity) of reprogramming for gene *i* is: R_*i*_ = αG_*i*_ (Fig. 3A). The total reprogramming (reprogramming expanse) for the set of genes that have to be upregulated is R_up_ = αΣG_*i*_ (Fig. 3B). The total reprogramming for the set of genes that have to be downregulated can be calculated similarly, but R_down_ is a negative value. The reprogramming constant α can be arbitrarily set as 1 for clarity, and the formulas become R_up_ = ΣG_*i*_ and R_down_ = *ΣG_*i*_* (Fig. 3C). The total amount of reprogramming (total reprogramming expanse) would be: R = R_up_ + |R_down_| (Fig. 3D). Based on this model, we have calculated the reprogramming expanse of human fibroblast reprogramming into pluripotency. The R_down_ is −11,096.4 LFC; R_up_ is 14,936.4 LFC, and the total reprogramming expanse R is 26,032.8 LFC (Table S3, and Fig. 3E). We also calculated the amount of reprogramming for the erasome and activatome to be −5,244.6 LFC and 10,190 LFC, respectively (Table S3). R_down_ is 74.3% of R_up_, while the amount of reprogramming for erasome is 51.5% that of activatome. These data indicate that upreprogramming is more dramatic than downreprogramming.

**Fig. 3.**
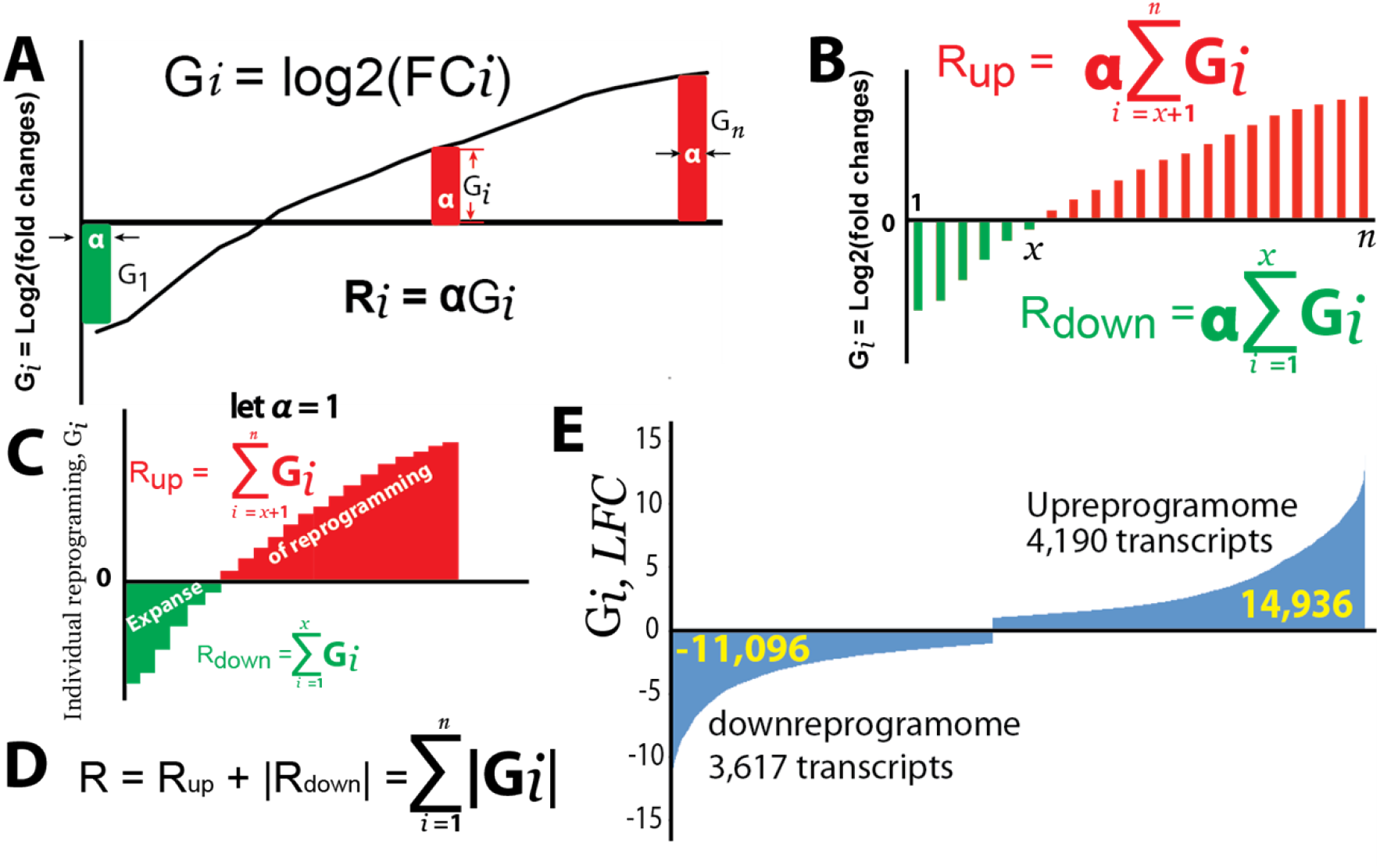
A mathematical model for quantification of reprogramming under an ideal condition (e.g., in a fertilized egg or a re-constructed egg with a somatic nucleus transferred into an enucleated oocyte). (A) A mathematical model for calculating reprogramming amount of an individual gene. (B) Mathematical models for calculating total reprogramming for downreprogramming (green) and upreprogramming (red). (C) A mathematical model for calculating the total reprogramming when an arbitrary reprogramming constant *a* is set to be 1. The x-axis is number of genes (from 1 to n) ordered based on log2(FC) values from low to high. (D) The formula for calculating the total reprogramming. (E) Waterfall plots showing amount of reprogramming of human fibroblast reprogramming into pluripotency.

Since WNT pathways are intensely and extensively reprogrammed, we also quantitate the amount of reprogramming in WNT pathways. The WNT downreprogramming R_WNT-down_ is −166.8 LFC while the R_WNT-up_ is 356.1 LFC (Fig. 2C). These results indicate that the components of WNT pathways are 2.1 times more upreprogrammed than downreprogrammed. The total WNT reprogramming R_WNT_ is 522.9 LFC, representing 2% of the overall reprogramming (Table S3).

### Profiling and quantification of reprogramming in cell morphorgenesis

To demonstrate further the utility of our quantification method for reprogramming, we analyzed genes involved in cellular morphogenesis and its regulation. Conversion of fibroblasts into iPSCs involves dramatic changes in cell morphology and establishes a unique cellular colony characteristic of PSC culture. In fact, high quality iPS cell lines can be established by selecting colonies with the characteristic cell and colony morphology without a reporter [7, 8, 20], and automatic imaging system can be used for identification of high quality iPSC colonies [21]. GO analyses of reprogramome reveal that 22 and 23 such GO terms are associated with the downreprogramome and upreprogramome, respectively (Table S6). Fig. 4B and 4D shows the top 10 GO terms under the category of cellular morphogenesis for upregulatome and downregulatome, respectively. There are 269 genes with roles in cellular morphogenesis that have to be downregulated at least 2 fold; and 252 such genes are in the upregulatome (Fig.s 4A, 4C, 4E). Thus, a total of 521 genes with roles in cellular morphogenesis have to be reprogrammed for pluripotency establishment, representing 3.8% of fibroblast transcriptome (Table S3 and S7). R_morph-down_ is −859.5 LFC while R_morph-up_ is 1,133 LFC, and the total reprogramming in cellular morphogenesis R_morph_ is 1,993 LFC, representing 7.7% of the overall reprogramming (Fig. 4E, and Table S3). These two data indicate that genes in cellular morphogenesis have to be reprogrammed more in intensity than extensiveness (7.7% vs 3.8%). That is, the average reprogramming of each genes in cellular morphogenesis (3.8 LFC, equivalent to 13.9 FC for each gene) is higher than that for the entire reprogramome (3.3 LFC, equivalent to 9.9 FC for each gene). Although there are more genes in cellular morphogenesis that have to be downregulated, the upreprogramming for such genes are more pronounced since R_morp-down_ is only 75.9% of R_morph-up_. In addition, the erasome includes 56 genes with roles in cellular morphogenesis while the activatome contains 130 genes with such roles (Table S7 and Fig. S2), indicating that upreprogramming plays a more critical role in the establishment of cell and colony morphology of iPSCs. In sum, there is intensive and extensive reprogramming in genes with roles in cellular morphogenesis.

**Fig. 4.**
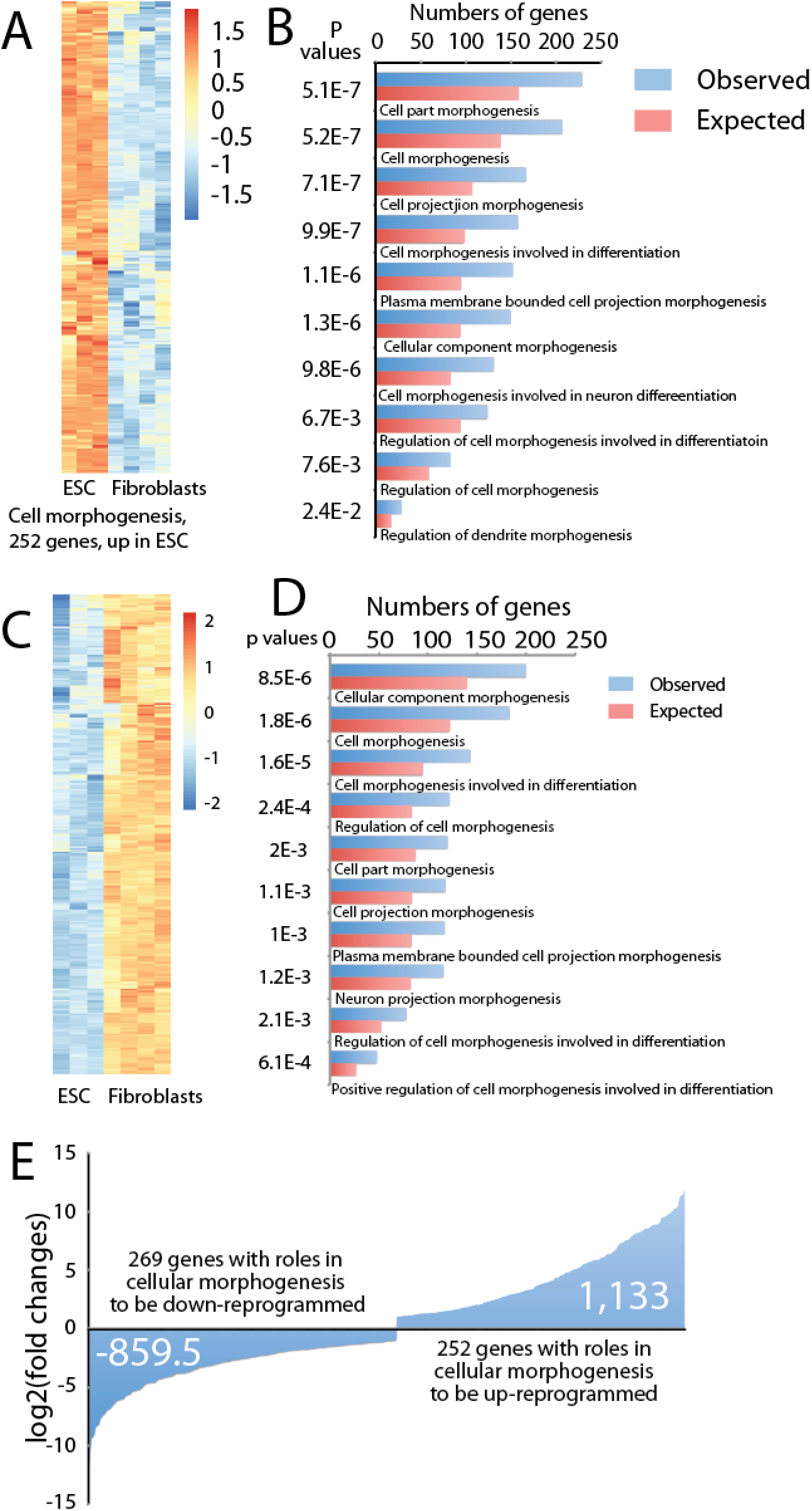
Gene sets in cellular morphogenesis that needs to be reprogrammed. (A) Heat maps showing genes regulating cellular morphogenesis that have to be upreprogrammed. (B) Top GO terms under the category of cellular morphogenesis for upreprogramome. (C) Heat maps showing genes regulating cellular morphogenesis that have to be downreprogrammed. (D) Top GO terms under the category of cellular morphogenesis for downreprogramome. (E) Quantification of reprogramming for genes with roles in cellular morphogenesis.

## Discussion

In this report, we have developed a new concept, reprogramome. This is analogous to transcriptome, epigenome, and kinome. It is more analogous to interferome [22]. This novel concept allows for fine profiling of genes to be reprogrammed. Further GO analyses, in combination with reprogramome profiling will provide many more insights into reprogramming. This is important because it is difficult to study the molecular mechanism because of very low efficiency of pluripotency reprogramming using the current protocols [5, 13]. With less than 1% of cells going to the pluripotent state, the signals we gain from the reprogramming population are mostly noise. For example, using our concept of reprogramome we were able to reveal that WNT pathways have to be extensively and intensively reprogrammed, and that the reprogramming is complicated although upreprogramming is predominant. This is not surprising because WNT pathways regulate various cellular and developmental processes. WNT pathways are complicated. There are canonical and at least two non-canonical WNT pathways, and these are interconnected. In the human genome there are 19 WNT ligands, more than 15 WNT receptors or co-receptors, and many downstream effectors [23]. WNT roles in reprogramming and pluripotency are poorly understood and warrant further investigation.

Another case study using reprogramome concept is the genes with roles in cellular morphogenesis and its regulation. We revealed that as high as 7.7% reprogramming involves genes that control and/or regulate cellular morphogenesis. This is also not surprising because iPSC generation involves dramatic changes in cellular morphology, and requires the establishment of strong cell-cell interaction among iPSCs and formation of a characteristic pluripotency colony. Our results are in agreement with a report that 1,454 genes were related to unusual colony morphology of human PSCs [24]. The reprogramome for genes in morphogenesis should be much greater because we focused on cellular morphogenesis and excluded genes for morphogenesis of tissues and organ in our current analyses.

With the concept of reprogramome, here we further developed a mathematic model to quantitate reprogramming. This also allows for quantification of the sub-reprogramomes including upreprogramome, downreprogramome, erasome, and activatome, as well as reprogramming for a specific pathway (e.g. WNT pathway), cellular functions or processes (e.g., cellular morphogenesis).

Our concept and methods can be applied to pluripotency reprogramming from other starting cell types, as well as reprogramming to various lineages such as neural and cardiac reprogramming. Furthermore, our concepts and methods can be applied to study the epigenetic changes required for a complete conversion of cell fates, i.e., epireprogramome. Of note, the same concept and methods may be applied to study the differences in transcription and epigenetics between any two types of cells including differences between cancer and their corresponding normal cells.

## Materials and methods

### Cell lines and tissue culture

We used and reported to our sponsors two NIH-registered human embryonic stem cell lines meeting federal and university regulations. We culture human embryonic stem cells (H1 and H9) in the chemically defined E8 media [25]. Human foreskin BJ fibroblasts (ATCC, CRL-2522) were cultured in fibroblast medium: DMEM, 10% heat-inactivated FBS, 0.1 mM 2-mercaptoethanol, 100 U ml^−1^ penicillin, 100 μg ml^−1^ streptomycin, 0.1 mM MEM NEAA and 4 ng ml^−1^ human bFGF.

### RNA preparation

Cells were harvested with Trizol reagent and stored at −80 °C until use. Total RNA was extracted using the Direct-zol Miniprep kit (Zymo Research, R2052). The four RNA samples of fibroblasts for RNA-seq were harvested on different days at different passage number. The three ESC RNA samples for RNA-seq were from two different ESC lines. For the repeat RNA samples of H1, they are harvested from different passages on different days.

### RNA-Seq

mRNA-sequencing was carried out on the Illumina HiSeq2500 following the established protocols. RNA-seq library preparation was done using the Agilent SureSelect Stranded kit (Agilent, Santa Clara, CA) as per the manufacturer’s instruction. The libraries were quantitated using qPCR in a Roche LightCycler 480 with the Kapa Biosystems kit for library quantitation (Kapa Biosystems, Woburn, MA) both immediately prior to and after library construction. We conducted paired end 50-bp sequencing for downstream analyses.

### Bioinformatics

All samples contained a minimum of 28.1 million reads with an average number of 40.1 million reads across all biological replicates. The FASTQ files were uploaded to the UAB High Performance Computer cluster for bioinformatics analysis with the following custom pipeline built in the Snakemake workflow system (v5.2.2) [26]: first, quality and control of the reads were assessed using FastQC, and trimming of the bases with quality scores of less than 20 was performed with Trim_Galore! (v0.4.5). All samples passed initial FASTQ QC, which included good quality scores through the read length and minimal adapter contamination. Following trimming, the transcripts were quasi-mapped and quantified with Salmon [27] (v0.12.0, with ‘--gencode’ and ‘-k 21’ flags for index generation and ‘-l A, ‘--gcBias’ and ‘-- validateMappings’ flags for quasi-mapping) to the hg38 human transcriptome from Gencode release 29. The average quasi-mapping rate was 88.8% and the logs of reports were summarized and visualized using MultiQC [28] (v1.6). The quantification results were imported into a local RStudio session (R version 3.5.3) and the package “tximport” [29] (v1.10.0) was utilized for gene-level summarization. Differential expression analysis was conducted with DESeq2 package [30](v1.22.1).

We prepared heat maps in RStudio using the package of *pheatmap* [31]; box plots with the package of ggplot2; ladder plots with the package of *plotrix*.

### Read count cutoff of DESeq2 data for expressed gene

We used two types of cutoffs for DESeq2 read counts for genes considered expressed, mean read count and individual read count cutoff. In DESeq2 normalization, we previously used a mean normalized read counts of 50 for the cell type in question as a cutoff for a gene to be considered active [7, 8]. This cutoff is supported by our current data (table S1). To confirm our selection of 50 as the mean-read-count cutoff in our data used in this manuscript, we manually selected three groups of genes based on our experience and previous microarray data [10], pluripotency (26 genes), fibroblast (21 genes), and double negative genes (20 genes), the last of which are known not to be expressed in both human fibroblasts and ESCs. For all these 67 genes except for POU5F1 (i.e., *OCT4)*, the mean normalized DESeq2 read counts range from 0 to 48.6 in the cell type in which they are not expressed. Most of these read counts are below 30, and only 2 of those are over 40. However, *OCT4* has a mean DESeq2 read counts of 124.6 in human fibroblasts. This is because we used FGF2 in our culture of fibroblasts. FGF2 was reported to stimulate expression of *OCT4* in fibroblasts [11]. The RNA-seq signals of OCT4 provide additional evidence that our RNA-seq is very sensitive and of high quality. In addition, many well-known pluripotency genes have read counts in the lower half of three-digit numbers, for examples, *DPPA2* (254), *GDF3* (272), *LEFTY2* (431), and *NODAL* (454). These read counts are in agreement with our previous microarray data, which displayed low levels of expression for these genes [10]. As an autocrine factor regulated by OCT4 and SOX2 in human ESCs with a role in ESC self-renewal [32], *FGF4* is considered a gene characteristic of hPSCs based on a survey of 59 human ESC lines from 17 laboratories by The International Stem Cell Initiative [33] because its expression strongly correlates with that of *NANOG.* But FGF4 expression level is very low [33] serving a reference gene for the lower limit. Our data with *FGF4* are in agreement with that of The International Stem Cell Initiative, and the averaged normalized mean read counts for *FGF4* for human ESCs are 80.6 versus 0.3 for fibroblasts. Therefore, the mean DESeq2 read counts of 50 is a reasonable cutoff (for example, this cutoff retains *FGF4* as an expressed gene but *CD19* and *CGB7* as inactive genes in human ESCs) (see table S1). To be stricter in selecting reliable expressed genes, we further used an individual-read-count cutoff of 10. That is, we further excluded genes from the list obtained using the above criteria, for which the individual normalized read count is less than 10 for any of the repeat experiments.

### Additional selection criteria

In addition to the read count cutoffs described above, we used other strict criteria to define the reliable reprogramomes. We use q values rather than p values. We used q values of <0.01 rather than <0.05. Furthermore, we included genes only with a least 2 fold of differences in expression levels rather than 1.5 fold as a cutoff.

## Supplementary Information

Supplemental information includes two Fig.s and seven tables.

## Author contributions

**Conceptualization**: Kejin Hu

**Data curation**: Kejin Hu, Lara Ianov, David Crossman

**Formal analysis**: Lara Ianov, Kejin Hu, David Crossman

**Investigation**: Kejin Hu

**Methodology**: Kejin Hu, Lara Ianov, David Crossman

**Project Administration**: Kejin Hu

**Visualization**: Lara Ianov, Kejin Hu

**Writing – original draft**: Kejin Hu

**Writing – review & editing**: Kejin Hu, Lara Ianov, David Crossman

## Funding

This research is supported by the National Institutes of Health (1R01GM127411) and the American Heart Association (17GRNT3367080).

## Acknowledgements

We appreciate the administrative support by Dr. Craig M. Powell, and technical help in RNA-sequencing by Dr. Michael Crowley. We also thank our colleagues at UAB for their critical reading of this manuscript.

## Declaration of Competing Interest

All authors declare no potential conflict of interests.

## Data availability statement

All raw RNA-seq data are being made accessible at the Gene Expression Omnibus (GEO) database repository with the access code of GSE138170 (available to public after publication). Analyzed data are within the article as figures and tables.

---

**Supplemental Fig. 1.**
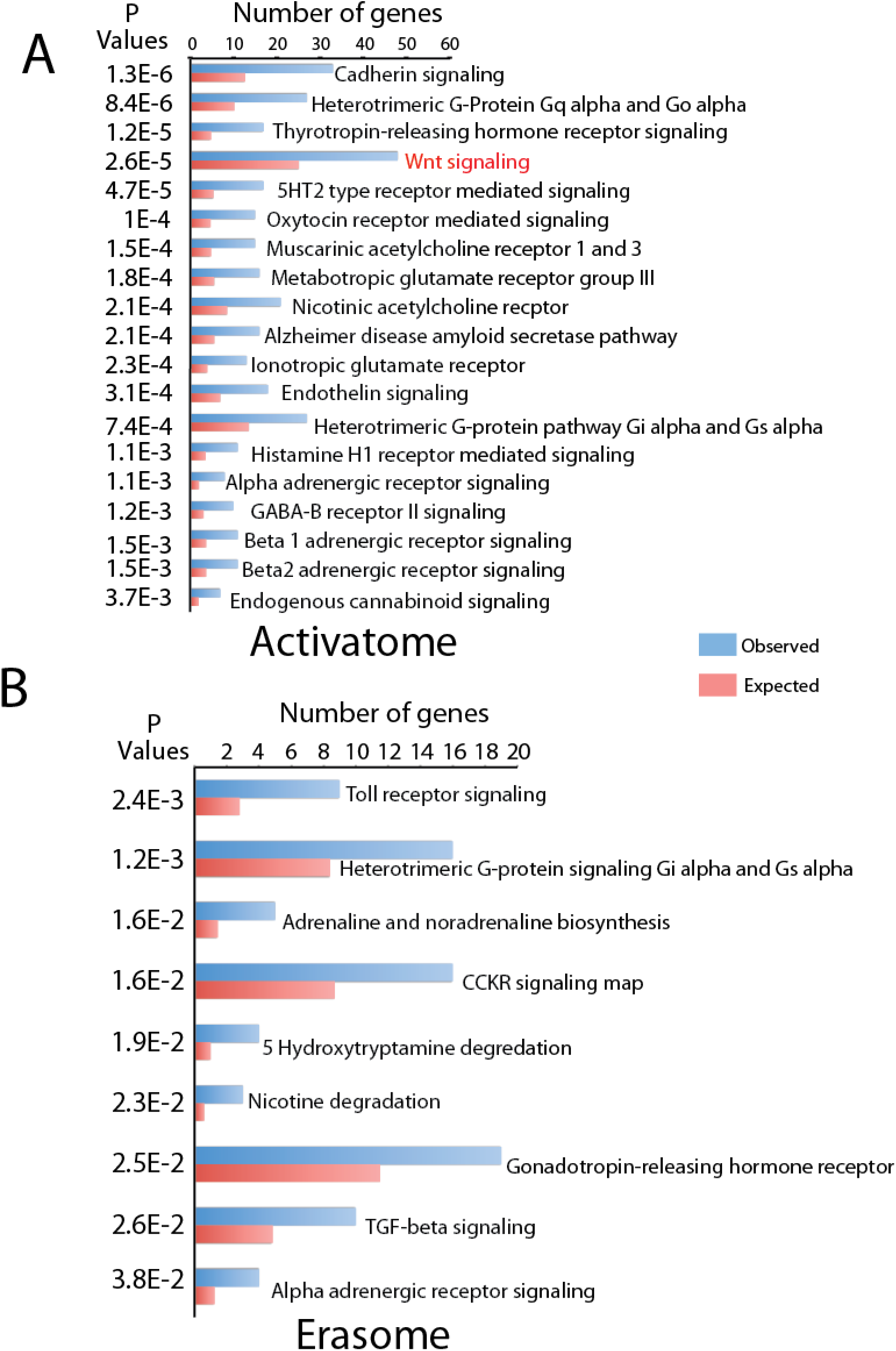
Top PANTHER pathway GO terms for activatome and erasome. (A) activatome. (B) erasome. Ranked by p values from low to high.

**Supplemental Fig. 2.**
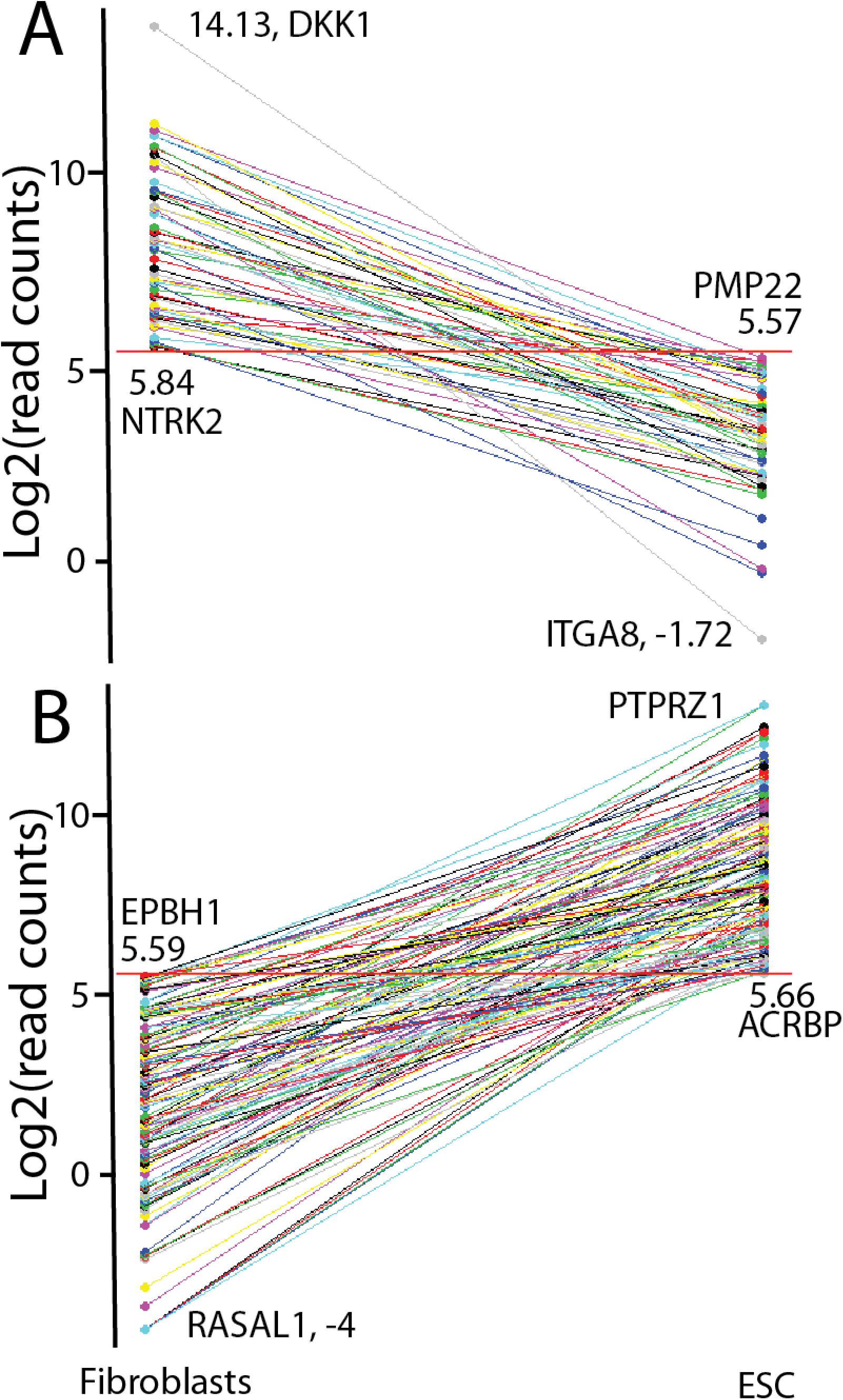
Ladder plots showing expression differences for genes with roles in cellular morphogenesis in the erasome (A) and activatome (B). The red horizontal lines indicate the log2(read counts) threshold for genes considered as being expressed. Labeled are genes with the lowest and highest read counts.

## References

[1] S. Markoulaki, A. Meissner, R. Jaenisch, Somatic cell nuclear transfer and derivation of embryonic stem cells in the mouse, Methods, 45 (2008) 101–114.

[2] J.A. Byrne, D.A. Pedersen, L.L. Clepper, M. Nelson, W.G. Sanger, S. Gokhale, D.P. Wolf, S.M. Mitalipov, Producing primate embryonic stem cells by somatic cell nuclear transfer, Nature, 450 (2007) 497–502.

[3] K. Takahashi, S. Yamanaka, Induction of pluripotent stem cells from mouse embryonic and adult fibroblast cultures by defined factors, Cell, 126 (2006) 663–676.

[4] J. Yu, M.A. Vodyanik, K. Smuga-Otto, J. Antosiewicz-Bourget, J.L. Frane, S. Tian, J. Nie, G.A. Jonsdottir, V. Ruotti, R. Stewart, Slukvin, II, J.A. Thomson, Induced pluripotent stem cell lines derived from human somatic cells, Science, 318 (2007) 1917–1920.

[5] K. Hu, All roads lead to induced pluripotent stem cells: the technologies of iPSC generation, Stem Cells Dev, 23 (2014) 1285–1300.

[6] K. Takahashi, K. Tanabe, M. Ohnuki, M. Narita, T. Ichisaka, K. Tomoda, S. Yamanaka, Induction of pluripotent stem cells from adult human fibroblasts by defined factors, Cell, 131 (2007) 861–872.

[7] Z. Shao, C. Yao, A. Khodadadi-Jamayran, W. Xu, T.M. Townes, M.R. Crowley, K. Hu, Reprogramming by De-bookmarking the Somatic Transcriptional Program through Targeting of BET Bromodomains, Cell Rep, 16 (2016) 3138–3145.

[8] Z. Shao, R. Zhang, A. Khodadadi-Jamayran, B. Chen, M.R. Crowley, M.A. Festok, D.K. Crossman, T.M. Townes, K. Hu, The acetyllysine reader BRD3R promotes human nuclear reprogramming and regulates mitosis, Nat Commun, 7 (2016) 10869.

[9] J.A. Thomson, J. Itskovitz-Eldor, S.S. Shapiro, M.A. Waknitz, J.J. Swiergiel, V.S. Marshall, J.M. Jones, Embryonic stem cell lines derived from human blastocysts, Science, 282 (1998) 1145–1147.

[10] K. Hu, I. Slukvin, Induction of Pluripotent Stem Cells from Umbilical Cord Blood, in: R.A. Meyers (Ed.) Reviews in Cell Biology and Molecular Medicine, Wiley - VCH Verlag GmbH & Co. KGaA, 2012, pp. 1–25.

[11] M. Jez, S. Ambady, O. Kashpur, A. Grella, C. Malcuit, L. Vilner, P. Rozman, T. Dominko, Expression and differentiation between OCT4A and its Pseudogenes in human ESCs and differentiated adult somatic cells, PLoS One, 9 (2014) e89546.

[12] L. Kang, C. Yao, A. Khodadadi-Jamayran, W. Xu, R. Zhang, N.S. Banerjee, C.W. Chang, L.T. Chow, T. Townes, K. Hu, The Universal 3D3 Antibody of Human PODXL Is Pluripotent Cytotoxic, and Identifies a Residual Population After Extended Differentiation of Pluripotent Stem Cells, Stem Cells Dev, 25 (2016) 556–568.

[13] K. Hu, Vectorology and factor delivery in induced pluripotent stem cell reprogramming, Stem Cells Dev, 23 (2014) 1301–1315.

[14] J. Jiang, Y.S. Chan, Y.H. Loh, J. Cai, G.Q. Tong, C.A. Lim, P. Robson, S. Zhong, H.H. Ng, A core Klf circuitry regulates self-renewal of embryonic stem cells, Nat Cell Biol, 10 (2008) 353–360.

[15] K.K. Chan, J. Zhang, N.Y. Chia, Y.S. Chan, H.S. Sim, K.S. Tan, S.K. Oh, H.H. Ng, A.B. Choo, KLF4 and PBX1 directly regulate NANOG expression in human embryonic stem cells, Stem Cells, 27 (2009) 2114–2125.

[16] A.M. Ghaleb, V.W. Yang, Kruppel-like factor 4 (KLF4): What we currently know, Gene, 611 (2017) 27–37.

[17] J.M. Shields, R.J. Christy, V.W. Yang, Identification and characterization of a gene encoding a gut-enriched Kruppel-like factor expressed during growth arrest, J Biol Chem, 271 (1996) 20009–20017.

[18] H. Mi, A. Muruganujan, J.T. Casagrande, P.D. Thomas, Large-scale gene function analysis with the PANTHER classification system, Nat Protoc, 8 (2013) 1551–1566.

[19] K. Hu, On Mammalian Totipotency: What Is the Molecular Underpinning for the Totipotency of Zygote?, Stem Cells Dev, 28 (2019) 897–906.

[20] K. Hu, J. Yu, K. Suknuntha, S. Tian, K. Montgomery, K.D. Choi, R. Stewart, J.A. Thomson, Slukvin, II, Efficient generation of transgene-free induced pluripotent stem cells from normal and neoplastic bone marrow and cord blood mononuclear cells, Blood, 117 (2011) e109–119.

[21] K. Tokunaga, N. Saitoh, I.G. Goldberg, C. Sakamoto, Y. Yasuda, Y. Yoshida, S. Yamanaka, M. Nakao, Computational image analysis of colony and nuclear morphology to evaluate human induced pluripotent stem cells, Sci Rep, 4 (2014) 6996.

[22] I. Rusinova, S. Forster, S. Yu, A. Kannan, M. Masse, H. Cumming, R. Chapman, P.J. Hertzog, Interferome v2.0: an updated database of annotated interferon-regulated genes, Nucleic Acids Res, 41 (2013) D1040–1046.

[23] C. Niehrs, The complex world of WNT receptor signalling, Nat Rev Mol Cell Biol, 13 (2012) 767–779.

[24] R. Kato, M. Matsumoto, H. Sasaki, R. Joto, M. Okada, Y. Ikeda, K. Kanie, M. Suga, M. Kinehara, K. Yanagihara, Y. Liu, K. Uchio-Yamada, T. Fukuda, H. Kii, T. Uozumi, H. Honda, Y. Kiyota, M.K. Furue, Parametric analysis of colony morphology of non-labelled live human pluripotent stem cells for cell quality control, Sci Rep, 6 (2016) 34009.

[25] G. Chen, D.R. Gulbranson, Z. Hou, J.M. Bolin, V. Ruotti, M.D. Probasco, K. Smuga-Otto, S.E. Howden, N.R. Diol, N.E. Propson, R. Wagner, G.O. Lee, J. Antosiewicz-Bourget, J.M. Teng, J.A. Thomson, Chemically defined conditions for human iPSC derivation and culture, Nat Methods, 8 (2011) 424–429.

[26] J. Koster, S. Rahmann, Snakemake--a scalable bioinformatics workflow engine, Bioinformatics, 28 (2012) 2520–2522.

[27] R. Patro, G. Duggal, M.I. Love, R.A. Irizarry, C. Kingsford, Salmon provides fast and bias-aware quantification of transcript expression, Nat Methods, 14 (2017) 417–419.

[28] P. Ewels, M. Magnusson, S. Lundin, M. Kaller, MultiQC: summarize analysis results for multiple tools and samples in a single report, Bioinformatics, 32 (2016) 3047–3048.

[29] C. Soneson, M.I. Love, M.D. Robinson, Differential analyses for RNA-seq: transcript-level estimates improve gene-level inferences, F1000Res, 4 (2015) 1521.

[30] M.I. Love, W. Huber, S. Anders, Moderated estimation of fold change and dispersion for RNA-seq data with DESeq2, Genome Biol, 15 (2014) 550.

[31] R. Kolde, pheatmap: pretty heatmaps, in, https://cran.r-project.org/web/packages/pheatmap/index.html, 2019.

[32] Y. Mayshar, E. Rom, I. Chumakov, A. Kronman, A. Yayon, N. Benvenisty, Fibroblast growth factor 4 and its novel splice isoform have opposing effects on the maintenance of human embryonic stem cell self-renewal, Stem Cells, 26 (2008) 767–774.

[33] I. International Stem Cell, O. Adewumi, B. Aflatoonian, L. Ahrlund-Richter, M. Amit, P.W. Andrews, G. Beighton, P.A. Bello, N. Benvenisty, L.S. Berry, S. Bevan, B. Blum, J. Brooking, K.G. Chen, A.B. Choo, G.A. Churchill, M. Corbel, I. Damjanov, J.S. Draper, P. Dvorak, K. Emanuelsson, R.A. Fleck, A. Ford, K. Gertow, M. Gertsenstein, P.J. Gokhale, R.S. Hamilton, A. Hampl, L.E. Healy, O. Hovatta, J. Hyllner, M.P. Imreh, J. Itskovitz-Eldor, J. Jackson, J.L. Johnson, M. Jones, K. Kee, B.L. King, B.B. Knowles, M. Lako, F. Lebrin, B.S. Mallon, D. Manning, Y. Mayshar, R.D. McKay, A.E. Michalska, M. Mikkola, M. Mileikovsky, S.L. Minger, H.D. Moore, C.L. Mummery, A. Nagy, N. Nakatsuji, C.M. O’Brien, S.K. Oh, C. Olsson, T. Otonkoski, K.Y. Park, R. Passier, H. Patel, M. Patel, R. Pedersen, M.F. Pera, M.S. Piekarczyk, R.A. Pera, B.E. Reubinoff, A.J. Robins, J. Rossant, P. Rugg-Gunn, T.C. Schulz, H. Semb, E.S. Sherrer, H. Siemen, G.N. Stacey, M. Stojkovic, H. Suemori, J. Szatkiewicz, T. Turetsky, T. Tuuri, S. van den Brink, K. Vintersten, S. Vuoristo, D. Ward, T.A. Weaver, L.A. Young, W. Zhang, Characterization of human embryonic stem cell lines by the International Stem Cell Initiative, Nat Biotechnol, 25 (2007) 803–816.

